# Acute knockdown of extracellular matrix protein Tinagl1 disrupts heart laterality and pronephric cilia in zebrafish embryonic development

**DOI:** 10.1101/2020.06.05.136747

**Authors:** Hannah Neiswender, Ellen K. LeMosy

**Affiliations:** Department of Cellular Biology and Anatomy, Medical College of Georgia, Augusta University

## Abstract

A highly-conserved extracellular matrix protein, Tinagl1, modulates Wnt, integrin, TGF-β, and EGF-R signaling *in vitro,* but its significance *in vivo* has remained in doubt. To bypass possible genetic compensation by an ortholog encoded exclusively in mammalian genomes, we examine Tinagl1 function in zebrafish embryos. In this model, *tinagl1* mRNA is detected in the developing spinal cord and pronephros. Acute knockdown using either CRISPR/Cas9 somatic mutagenesis or splice-blocking morpholinos reveals left-right (LR) heart looping defects, pronephros dilatations, and ventral body curvature. This constellation of defects characteristically results from the loss of motile cilia function, and we confirm the presence of shortened and fewer cilia in the pronephric duct and in the Kupffer’s vesicle where LR asymmetry is established. A link to known Wnt3a/β-catenin signaling that activates the motile cilia transcriptional program is supported by manipulation of Wnt3a and β-catenin levels in *tinagl1* knockdown embryos. In addition to ciliopathy-like defects, the *tinagl1* knockdown shows disorganization of longitudinal axon tracts in the spinal cord and defects in motor neuron outgrowth. Together, these results provide evidence that Tinagl1 is important in development, and that zebrafish is an ideal model in which to explore its relationships to cilia and secreted signaling molecules.

## Introduction

The extracellular matrix (ECM) provides an information-rich structural environment for development and maintenance of tissues. While a subset of ECM components has been extensively studied, others that likely warrant study have been largely ignored (Stoeger et al., 2018). Among the latter group are the structurally related proteins Tinag (tubulo-interstitial nephritis antigen) and Tinagl1 (tubulo-interstitial nephritis antigen-like protein 1; also called TIN-ag-RP, LCN7, AZ-1, Arg-1) (Kobayashi et al., 2001; Mukai et al., 2003; Wex et al., 2001; Zhou et al., 2000). Each contains a somatomedin B domain, a lipocalin motif, and an apparently inactive cathepsin B-like domain in which a catalytic cysteine has been replaced by a serine residue (see **Supplemental Figure 1** and its legend for further information on these motifs). Studies *in vitro* have demonstrated physical interactions of *Drosophila* Tinagl1 with the Wg (Wnt family) protein, having positive cofactor activity in long-range signaling (Mulligan et al., 2012). Mammalian Tinagl1 physically interacts with integrins, blocking a ligand-binding site for fibronectin, and with the EGF receptor, blocking its dimerization and activation after EGF engagement (Shen et al., 2019). Together with conservation of orthologs throughout Bilaterian animal species and, for human Tinagl1, lower than predicted incidence of loss of function alleles (observed/expected ratio 0.38 [0.24-0.64] in gnomAD; Karczewski et al., 2020), these physical interactions with key signaling components suggest that this small gene family may have important but unrecognized roles in development and disease.

In adult mammals, Tinagl1 is expressed at varying levels in most epithelia and most prominently in vascular endothelium, but is also expressed by non-epithelial vascular smooth muscle and adrenal cortex (Mukai et al., 2003; Wex et al., 2001). Tinag expression is largely limited to the kidney (Ikeda et al., 2000). The proteins are localized to the basement membrane in epithelia and vascular smooth muscle, while Tinagl1 is also diffusely distributed in the glomerular interstitial matrix and in a gradient across the depth of the adrenal cortex (Charonis et al., 1994; Lennon et al., 2014; Mukai et al., 2003; Wex et al., 2001). Both proteins have been reported to mediate integrin-dependent adhesion of different cell types *in vitro,* and to potentiate cell adhesion onto structural matrix proteins including fibronectin (Chen et al., 1996; Kalfa et al., 1994; Li et al., 2007). However, Tinagl1 was recently demonstrated to inhibit integrin/FAK signaling in cancer cells (Shen et al., 2019). These results suggest that there may be context-, cell-type-, and family member-specific functions for Tinag and Tinagl1.

Exploration of these proteins in developmental contexts has been limited. Inhibition of Tinag expression in a mouse embryonic kidney explant model reduced tubulogenesis and ureteric branching morphogenesis, while glomerular differentiation was preserved (Kanwar et al., 1999). A Tinagl1 knockout mouse is viable without gross phenotypes, but has impaired female fertility with fetal losses across gestation (Takahashi et al., 2016). Although the investigators postulated that the placenta, normally enriched in Tinagl1 (Tajiri et al., 2010), is defective, the fetal defects were not explored. Thus, the role of Tinagl1 during development remains unclear.

We have established a zebrafish model in which the developmental roles of Tinagl1 can be more readily explored *in vivo*. Zebrafish demonstrate well-characterized, rapid, and easily visualized development of organ systems similar to those in mammals. Additionally, zebrafish have a single family member, Tinagl1, as observed in other non-mammalian model organism genomes. Here we describe expression of zebrafish *tinagl1* at time points across the first two days of embryogenesis, demonstrating dynamic, transient expression during early zebrafish development. We use morpholinos and G0 CRISPR/Cas9 somatic mutagenesis to acutely knock down Tinagl function, where we observed but did not explore the mildly delayed vasculogenesis of branches of the aorta previously reported to result from morpholino-mediated knockdown (Brown et al., 2010). Instead, we provide initial evidence for Tinagl1 functions in heart left-right patterning and in pronephros and nervous system development. These studies point toward there being important roles for Tinagl1 in vertebrate development.

## Materials and Methods

### Zebrafish Handling

The Tübingen wild type strain was maintained under standard conditions at 28.5°C in the Zebrafish Core Facility of Augusta University, which maintains AAALAC accreditation. All animal experiments were performed with approval and in accordance with guidelines of the Augusta University Institutional Animal Care and Use Committee [AUP 2013-0592]. Fish embryos were staged by developmental morphology (Kimmel et al., 1995).

### RNA *in situ* hybridization

Whole mount *in situ* hybridizations were performed using a standard protocol (Thisse and Thisse, 2008) with *tinagl1* (Neiswender et al., 2017) and *myl7* (Chen et al., 2008) probes.

### Acute Knockdown using Morpholinos

Splice-blocking *tinagl1* morpholinos and 5-bp mismatch control morpholinos were designed and synthesized by Gene Tools, LLC (Philomath, OR), as previously described (Neiswender et al., 2017). The Wnt3a MO has been previously described (Shimizu et al., 2005). Morpholinos were diluted in injection buffer (0.25% phenol red, 120 mM KCl, 20 mM Hepes-NaOH, pH 7.5), and 1 nl volume was injected into one-cell stage embryos. Final injected amounts were 3.5 – 10 ng of TEx1 MO or MM1,20 ng of TEx2 MO or MM2, and 3.5 – 8.5 ng of Wnt3a MO. Capped full-length *tinagl1, ctnnb2,* and *eGFP* mRNAs used in rescue experiments were synthesized from plasmid vectors using Ambion mMessage mMachine kits (Ambion/Life Technologies, Grand Island, NY). The purified mRNAs were quantified and 100 pg mRNA (amounts optimized for *tinagl1* and *ctnnb2* mRNA rescue) was combined with TEx1 MO prior to injection into one-cell stage embryos.

### Acute Knockdown using CRISPR/Cas9 Somatic Mutagenesis

The pCS2-nCas9n plasmid was a gift from Wenbiao Chen (Jao et al., 2013; Addgene plasmid #47929; http://n2t.net/addgene:47929; RRID:Addgene_47929). *Cas9* mRNA was generated using the Ambion mMessage mMachine SP6 kit following Not I digest and purification of the plasmid. Synthetic DNA templates for sgRNA synthesis were made using a published protocol (Gagnon et al., 2014). The constant oligonucleotide was annealed with genespecific oligonucleotides to *golden (slc24a5)* ATTTAGGTGACACTATA**GGTCTCTCGCAGGATGTTGC**GTTTTAGAGCTAGAAATAGCAAG (Jao et al., 2013) or *tinagl1* exon 2 ATTTAGGTGACACTATA**GGCTCTTACTGTCAGAGGAG**GTTTTAGAGCTAGAAATAGCAAG GGCTCTTACTGTCAGAGGAG identified using the CHOPCHOP algorithm v1.0 (Montague et al., 2014). Following fill-in with T4 DNA polymerase, Ambion MegaScript SP6 kit was used to generate the sgRNAs. One-cell embryos were injected with 1 nl of a solution containing 300 ng/μl sgRNA and 500 ng/μl Cas9 mRNA. These doses gave >75% embryo survival at 1 dpf, while providing at least 30% eye pigmentation mosaics among the *golden* sgRNA-injected embryos, and approximately 30% (10/29) *tinagl1* sgRNA-injected zebrafish demonstrating germline transmission to F1 embryos using T7E1 analysis of F1 embryos.

### Immunostaining and analysis of cilia

Primary antibodies used included anti-acetylated tubulin monoclonal 6-11B-1 (Sigma; 1:500 dilution) and primary motor neuron axon monoclonal antibody znp-1 deposited to the DSHB by B. Trevarrow (DSHB hybridoma product znp-1 supernatant; 1:5 dilution). Secondary antibodies were from Jackson ImmunoResearch. Imaging was performed on an upright Zeiss LSM 510 Meta confocal microscope in the Cell Imaging Core Laboratory based in the Cellular Biology and Anatomy Department at Augusta University. Cilia length was measured from z-stack data using length tools of the LSM 510 software. The anterior region of one pronephros per embryo was selected for counting cilia numbers per 100 μm length, and the lengths of 20 cilia from each of 20-30 embryos of each genotype were measured per experimental trial. For KV cilia, total cilia per KV were counted, and the lengths of 20 cilia from each of 30 embryos of each genotype were measured. Mean and standard deviation were calculated for combined cilia in each trial group. ANOVA was calculated and was < 0.001; pairwise comparisons were performed with Student’s t test to calculate *p*-values.

## Results

### Expression of *tinagl1* mRNA in zebrafish embryos

To identify tissues in which *tinagl1* function may be required in early development, we performed *in situ* mRNA hybridization of wild-type zebrafish embryos at time points across the first 48 hours of development (**Figure 1**). At the earlier times examined, *tinagl1* expression was detected in the nervous system (16 hpf, 22 hpf; **Figure 1B, D**). Expression was especially prominent in transverse bands aligned with the hindbrain, although in a relatively ventral position, and a pattern of scattered cells along the developing spinal cord that shifted caudally as development progressed. Floor plate expression could also be detected at 22-27 hpf (**Figure 1D**). By 35 hpf, only a few cells within the nervous system appeared to be stained (asterisks in **Figure 1E, G**).

**Figure 1.**
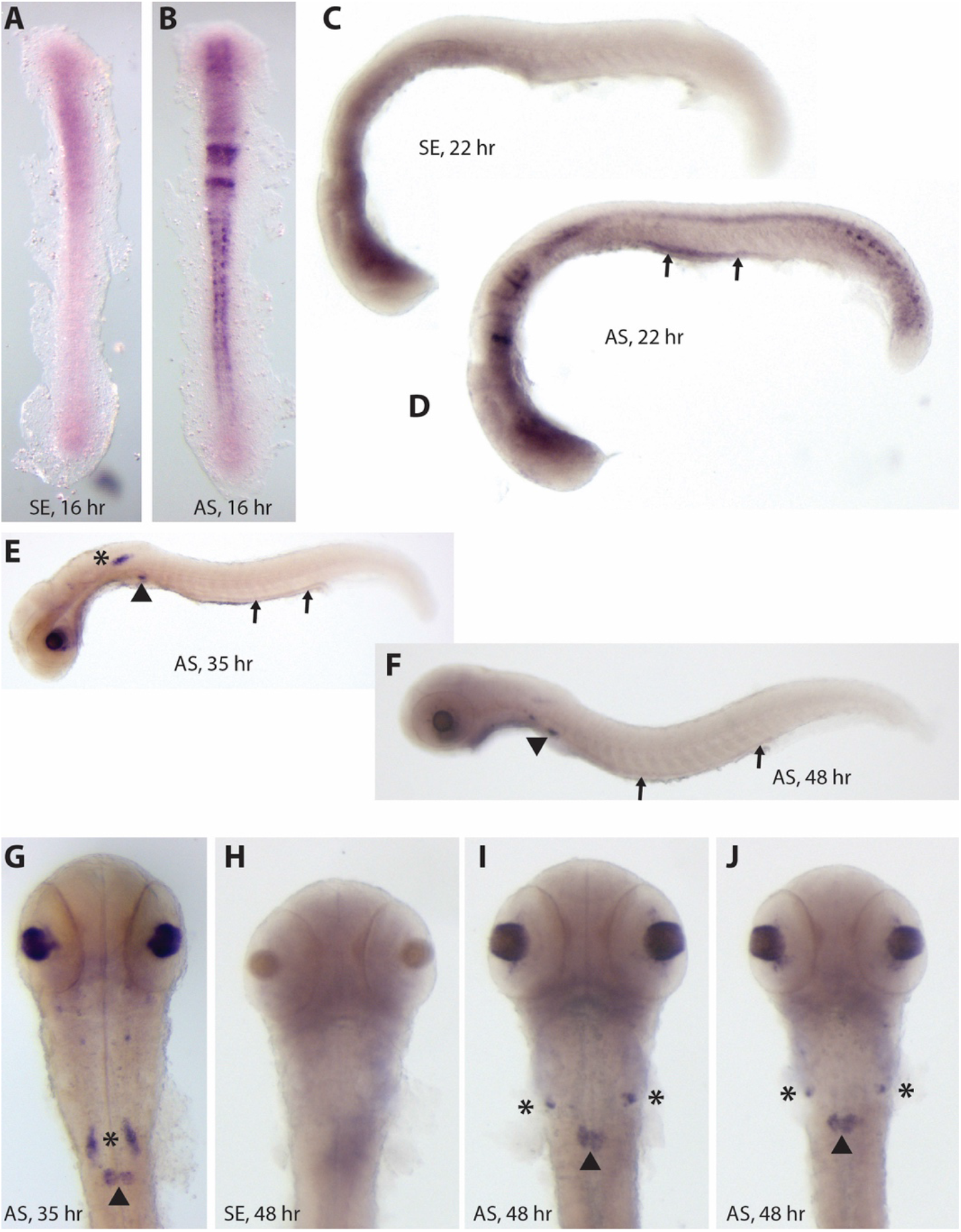
*tinagl1* mRNA expression in early development. Fixed embryos were hybridized with sense (SE; panels A, C, H) or antisense (AS; panels B, D, E-G, I, J) digoxygenin-labeled RNA probes. Time points were 16 hours (panels A, B), 22 hours (panels C, D), 35 hours (panels E, G), and 48 hours (panels F, H-J) post fertilization. Paired small arrows mark the pronephric duct, arrowheads mark the renal glomerulus, and asterisks mark a pair of unidentified cells that move ventrally, anteriorly, and laterally from 35 hpf to 48 hpf. Head is oriented to the left in lateral views (panels C-F), and to top in dorsal (panels A, B) or ventral (panels G-J) views.

The developing renal system also showed *tinagl1* expression. The pronephric duct showed strongest signal at 22-24 hpf, which diminished by 35 hpf and was nearly absent in 48 hpf embryos (**Figure 1D, F**). Renal glomerulus staining was seen at the later time points (**Figure 1E-J**). Together, these results suggest the spinal cord and pronephros are tissues in which zebrafish *tinagl1* could function, but do not rule out its action elsewhere, e.g., involving low expression not detected here or the diffusion of Tinagl1 once secreted.

### Two approaches demonstrate developmental defects when *tinagl1* expression is reduced

To determine whether Tinagl1 is required for early development, we first injected onecell zebrafish embryos with either of two morpholinos targeting splicing after the first or second coding *tinagl1* exons (**Figure 2B, C**, and **Supplemental Figures 2 and 3**). These morphant embryos displayed a common pattern of developmental defects including ventral body curvature, reduced eye size, frequent renal cysts, heart defects, and pericardial edema later associated with more generalized edema. Hydrocephalus was observed but not routinely scored. These defects were observed in parallel with craniofacial cartilage defects that we previously described (Neiswender *et al.,* 2017). Co-injection of *tinagl1* mRNA with the MOs resulted in partial to nearly-complete rescue of these phenotypes (**Figure 2D, E**; Neiswender *et al.,* 2017), and 5-bp mismatch (MM) morpholino-injected embryos demonstrated low to no defects depending on the defect scored (**Supplemental Figures 2 and 3**; Neiswender *et al.,* 2017). Co-injection of sub-threshold doses of TEx1 MO and TEx2 MO led to the same constellation of defects as either MO alone at higher doses (**Supplemental Figure 4**). The use of morpholinos affecting distinct splice sites, mRNA rescue, and absence of phenotypes with the corresponding MM morpholinos, all support the likelihood that these defects are due to specific loss of Tinagl1 (Eisen and Smith, 2008; Stainier et al., 2017).

**Figure 2.**
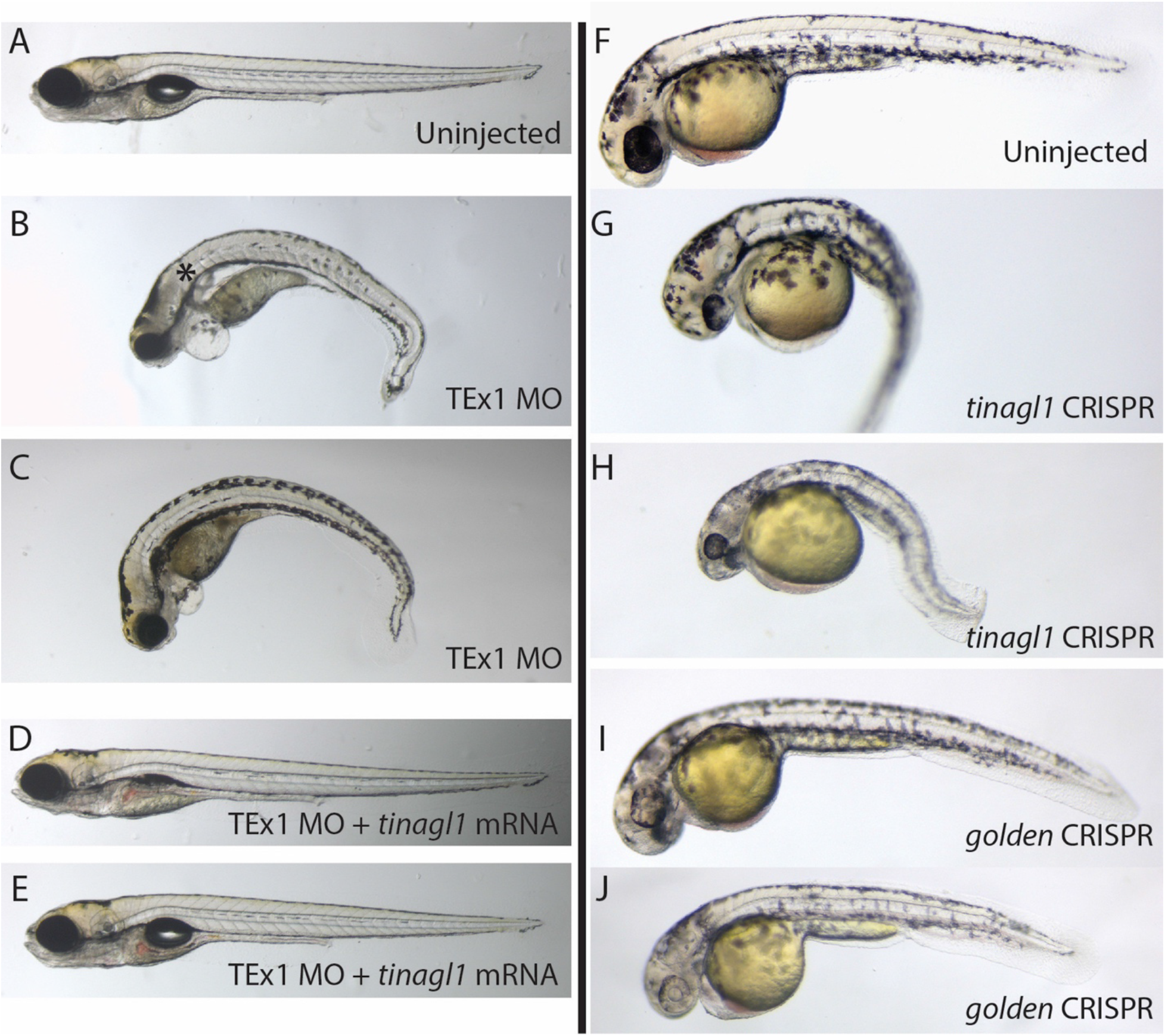
Gross phenotypes of *tinagl1* morphants and crispants. (A-E) Comparison of 5 dpf larvae that were either uninjected (normal; A), injected with TEx1 MO (B, C), or injected with TEx1 MO + 100 pg *tinagl1* mRNA (D, E). Asterisk in B is adjacent to a renal cyst. Embryo in E shows more complete rescue of the jaw compared to embryo in D. (F-J) Comparison of 2 dpf embryos that were either uninjected (F), injected with *tinagl1* sgRNA + Cas9 mRNA (G, H), or injected with *golden* sgRNA + Cas9 mRNA. Phenotypes are described in the main text.

As a second and independent approach to reduce *tinagl1* expression, we generated somatic mosaic mutant embryos (“crispants”) using CRISPR/Cas9 mutagenesis (Gagnon et al., 2014; Jao et al., 2013). Targeting the first coding exon of *tinagl1* resulted in a significant proportion of crispant embryos (~30%) that demonstrated ventral body curvature, reduced eye size, heart defects, and at a lower rate demonstrated craniofacial defects and renal cysts (**Figure 2G, H**). None of these defects were observed in embryos where the pigmentation gene *golden* was targeted as a general negative control for this mutagenesis method; instead, they showed patchy depigmentation as expected for *golden* crispants (**Figure 2I, J**). The similarity of phenotypes observed in both morphant and crispant embryos suggests that we are observing genuine requirements for *tinagl1* in development, versus technical artifacts.

### Heart laterality is altered in a significant proportion of *tinagl1* knockdown embryos

The heart appeared to be mispositioned in a proportion of *tinagl1* morphant and crispant embryos. To assess the nature of the heart defects more clearly, we used *in situ* hybridization with a *myl7* (cardiac myosin light chain) probe to label the hearts of 48 hpf embryos (**Figure 3**). Uninjected embryos at this age show heart tubes in which the future ventricle (dark purple in **Figure 3A**) is in a relatively rostral position and to the right of midline, while the atrium (light purple in **Figure 3A**) is more caudal and to the left of midline. Normal bending of the ventricular segment to the right between 30-48 hpf is called D-looping, while defects in earlier left-right (LR) asymmetry establishment or in heart tube migrations can result in reversed L-looping or in unlooped midline heart tubes (Bakkers, 2011). The *tinagl1* morphant embryos showed randomization of looping, with D-looped, L-looped, and midline unlooped hearts detected in broadly similar proportions (representative images in **Figure 3B-D**; scoring in **Figure 3I**). TEx1 MO produced an additional class that we scored as “no atrium” because of the apparent absence of the lighter-staining component representing this chamber (**Figure 3F**). These heart tubes were typically in the midline and in a rostral position within the chest, and appear to represent a more severe anomaly of heart development. The mismatch control morpholinos MM1 and MM2 showed some background of L-looping and “no atrium,” or of L-looping only, respectively, but did not approach the proportion of looping defects seen with the *tinagl1-* specific morpholinos (**Figure 3E, I**). To confirm these findings, we performed *myl7* staining on our *tinagl1* and *golden* crispants. The *tinagl1* crispants yielded ~27% L-looped hearts, while a negative control *golden* sgRNA yielded <5% L-looped hearts (**Figure 3G-I**). Midline and no atrium phenotypes were not observed.

**Figure 3.**
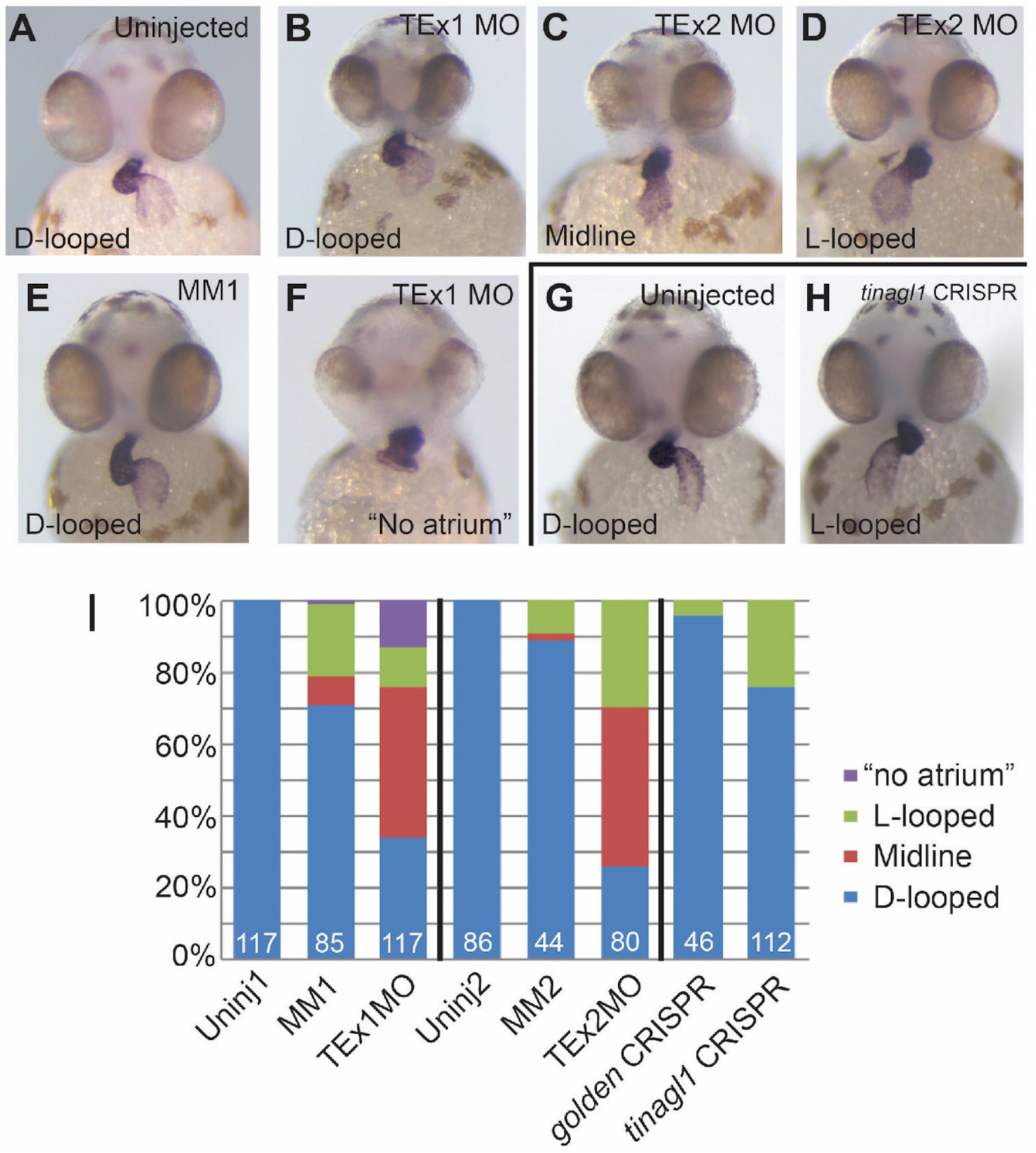
Laterality of heart looping is altered by *tinagl1* knockdown. Laterality was scored using a *myl7* antisense RNA probe that labels cardiac muscle. (A-H) Representative images of the phenotypes seen, including D-looped (normal), L-looped, midline unlooped, and a variant of midline heart tubes scored as “no atrium” due to absence of the lighter-staining caudal portion of the tube. Panels A-F show embryos from the morpholino experiments, and Panels G-H show embryos from the G0 CRISPR knockdown experiments. Panel I is a graphical comparison of the scored phenotypes. Numbers of embryos scored for each condition are indicated at the base of each bar, and conditions tested in parallel embryo pools are divided by vertical lines, e.g., TEx1 MO, its MM1 control, and Uninj1 were compared in same experiments.

Together these results show that both methods of acutely reducing *tinagl1* expression disrupt the normal lateralization of heart looping, suggesting Tinagl1 has a function in left-right (LR) patterning of the embryo. LR patterning is established by the activity of motile cilia. Intriguingly, many of the phenotypes observed in *tinagl1* morphants and crispants – heart looping defects, ventral body curvature, hydrocephalus, small eyes, and renal cysts – are suggestive of cilia defects (Essner et al., 2005; Kramer-Zucker et al., 2005). With this in mind, we next examined cilia in our models.

### Motile cilia are fewer and shorter in the Kupffer’s vesicle with MO knockdown of *tinagl1*

In vertebrates, LR patterning is established by the directional cilia-driven flow of fluid in the laterality organ. In zebrafish, this laterality organ is Kupffer’s vesicle (KV), a transient structure lined with motile cilia (Essner et al., 2005; Kramer-Zucker et al., 2005). If these cilia are disrupted, then the LR axis is not consistently established, leading to randomization of the laterality of organs including the heart. To test whether defective KV cilia could underlie the altered randomization of heart laterality in *tinagl1* knockdown embryos, we stained KV cilia for acetylated tubulin and examined their morphology in 15-16 hpf embryos. The KV cilia of MM1- or MM2-morpholino-injected embryos appeared similar to those of wild-type uninjected embryos, while KV cilia in TEx1 MO- or TEx2 MO-injected embryos were grossly disrupted (**Figure 4A-D**). Quantification showed reduction of cilium length by an average of 53-58% (**Figure 4E**), and reduction of cilia number per KV by an average of 30-37% (**Figure 4F**). Since reduction of either length or number of KV cilia can result in reduced fluid flow and failure to consistently establish the left, we hypothesize that dysmorphic cilia underlie the heart laterality defects in *tinagl1* knockdown embryos.

**Figure 4.**
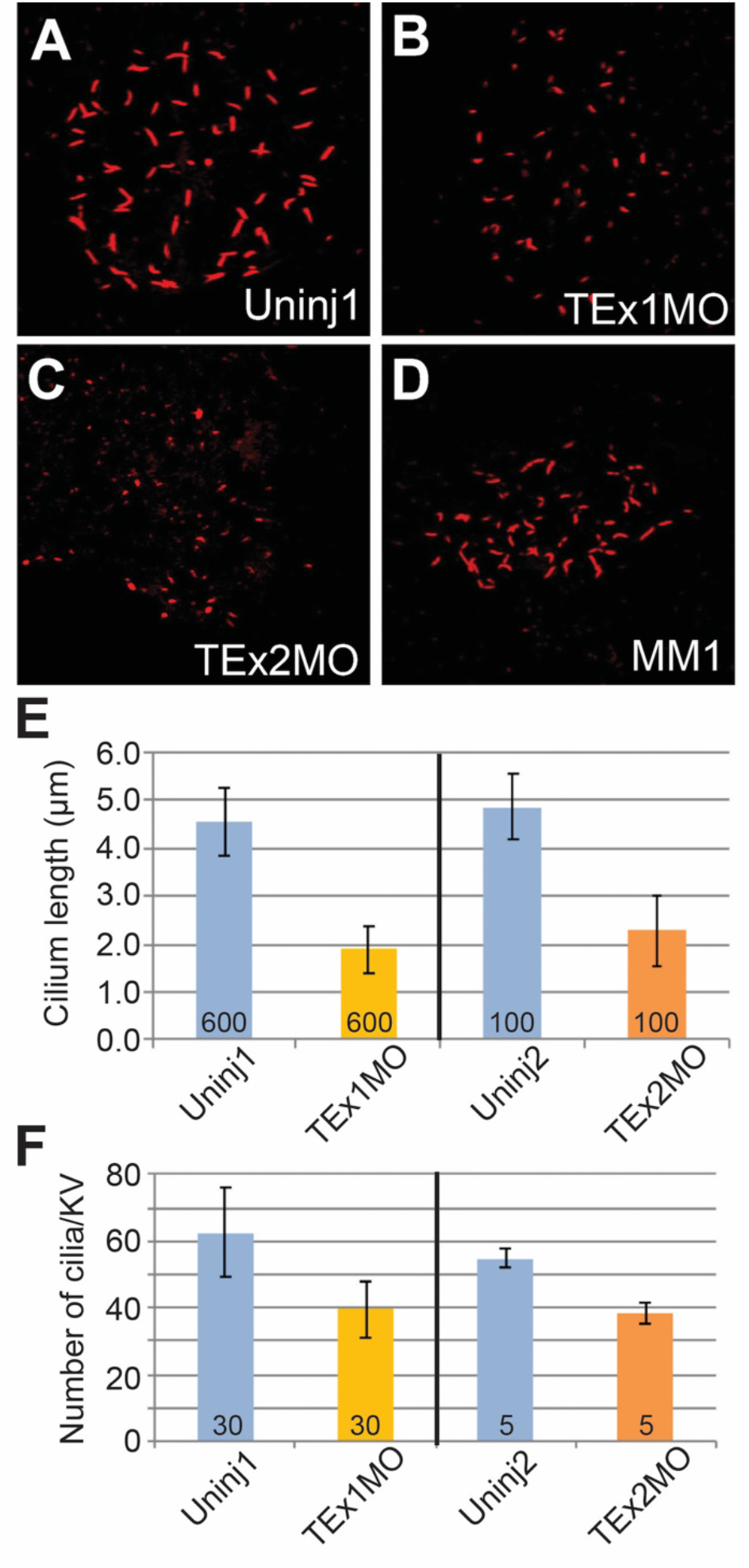
Cilia length and number are reduced in the Kupffer’s vesicle (KV) with *tinagl1* knockdown. KV cilia of 15-16 hpf fixed embryos were stained with anti-acetylated tubulin antibody. Panels A-D show representative Kupffer’s vesicles of uninjected (A), TEx1 MO and TEx2 MO-injected (B, C), and MM1-injected (D) embryos. In Panels E and F, cilium length (E) and cilium number per KV (F) were measured, respectively, with numbers at base of bars representing total number of cilia (E) or KVs (F) measured. Bars indicate s.d.

### Pronephric cilia are less organized, fewer, and shorter in *tinagl1* knockdown embryos

Motile cilia are similarly important in driving fluid flow in the renal system. Considering the strong expression of *tinagl1* in the pronephric duct, coupled with the prevalence of renal cysts upon *tinagl1* knockdown, we postulated that cilia morphology may be disrupted in this system. To test this hypothesis, we stained pronephric cilia for acetylated tubulin. We compared cilia in the anterior half of the 24 hpf pronephric duct in uninjected embryos and in embryos injected with TEx1 MO or its control MM1 morpholino. The overall organization of cilia was disrupted in TEx1 MO embryos, but not MM1 embryos (**Figure 5A**). Quantification showed reduction of cilium length by ~40% (**Figure 5B**) and reduction of cilia number per 100 μm anterior duct of ~50% (**Figure 5C**). Examination of G0 CRISPR/Cas9 knockdown embryos demonstrated similar disorganization in a proportion of *tinagl1* crispant embryos but not those treated with negative control *golden* sgRNA (**Figure 5D**); the length and number of cilia were not quantified for the CRISPR knockdown.

**Figure 5.**
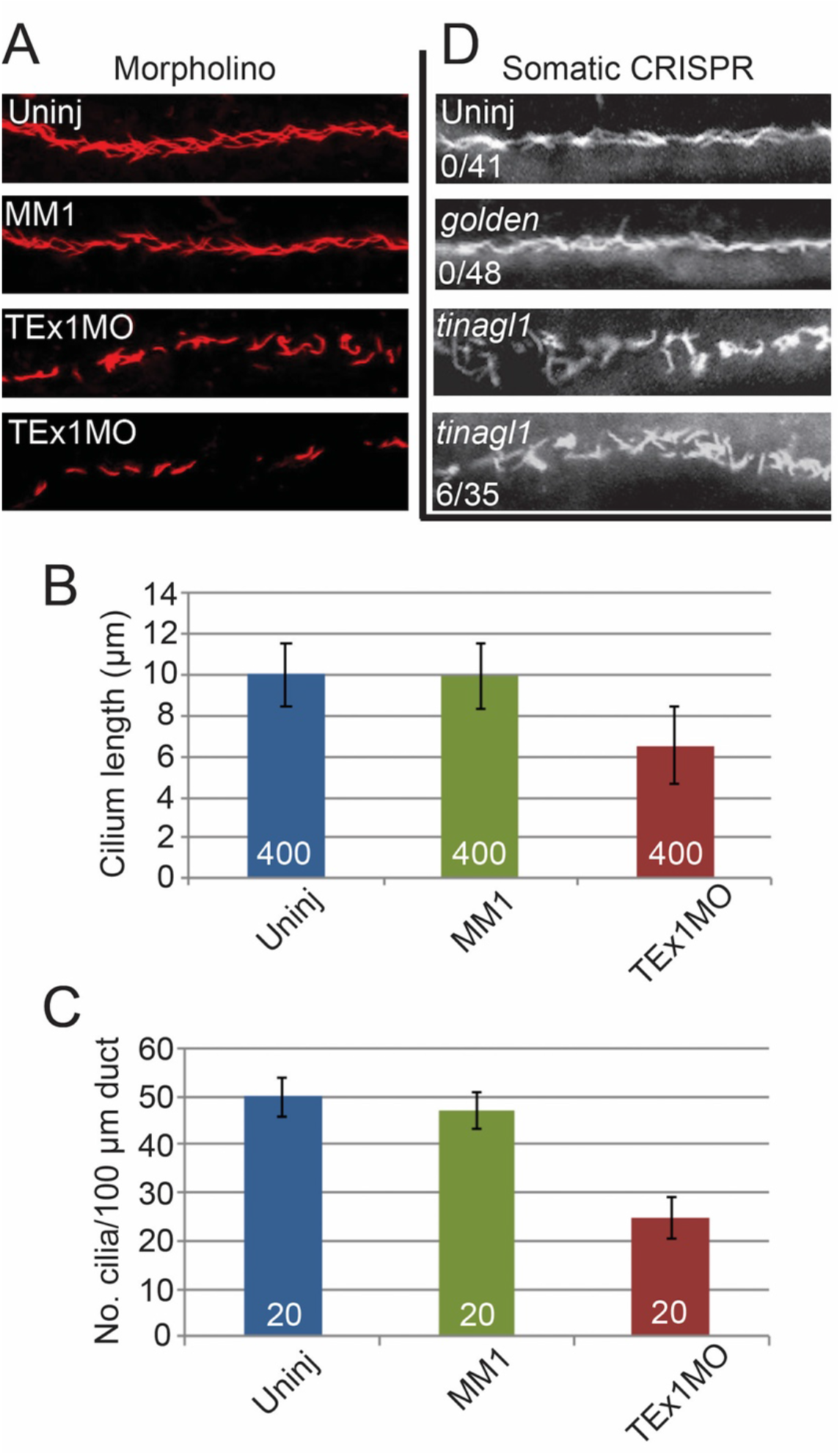
Acute *tinagl1* knockdown disrupts pronephric duct cilia. (A-C) Comparison of pronephric duct cilia appearance (A), cilium length (B), and cilia number per 100 μm length of pronephric duct (C) for uninjected, mismatch MM1-injected, and Tex1 MO-injected embryos. Total number of cilia (B) or embryos where a single pronephric duct per embryo was measured (C) are indicated by numbers at base of each bar. (D) Comparison of somatic G0 CRISPR embryos treated with *golden* or *tinagl1* guide RNAs, with numbers indicating embryos showing abnormal pronephric duct cilia out of total embryos stained and examined.

Consistent with our prior observations regarding partial rescue by *tinagl1* mRNA of craniofacial defects (Neiswender et al., 2017) and of ventral body curvature (**Figure 2**), we observed partial rescue of pronephric cilia length, number, and organization by *tinagl1* mRNA co-injected with TEx1 MO (**Figure 6A, D, E**) but not by co-injection of the same quantity of a control *eGFP* mRNA (**Figure 6C**).

**Figure 6.**
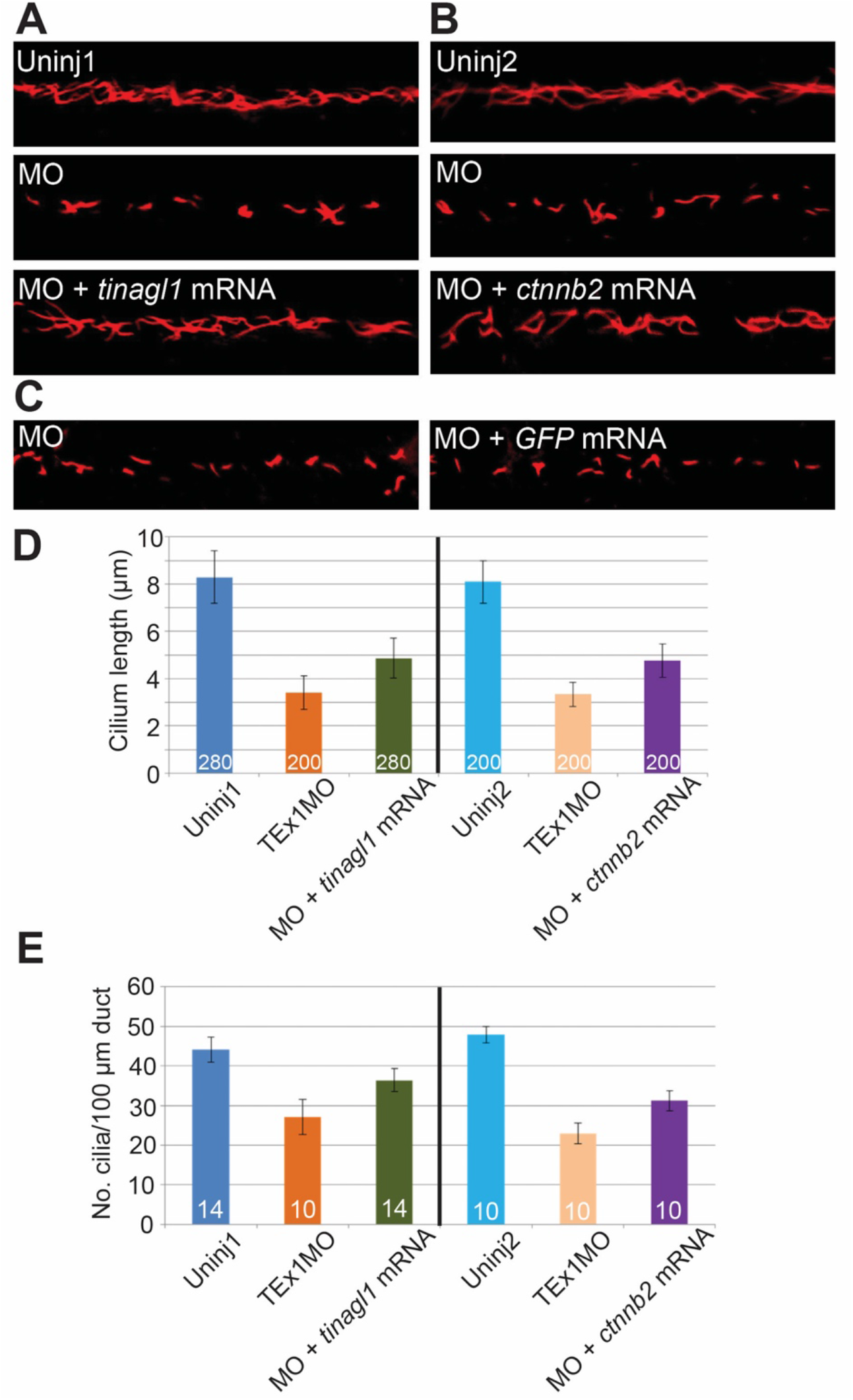
Pronephric duct cilia defects are partially rescued by *tinagl1* or *β-catenin (cttnb2)* mRNAs. (A-C) Representative images of pronephric duct cilia immunostaining at 24 hpf after injection of the indicated morpholinos alone or in combination with 100 pg/embryo *tinagl1* (A), *ctnnb2* (B), or control *GFP* (C) mRNA. (D) Quantitation of cilium length. (E) Quantitation of number of cilia per 100 μm length of pronephric duct. Labeling conventions are as in prior figures.

### *tinagl1* MO knockdown appears to interact with *Wnt3a* and *β-catenin* in regulation of pronephric cilia

Published studies provide support for the possibility that Tinagl1 could act in the formation or maintenance of cilia through promotion of Wnt signaling. The Nusse lab demonstrated a strong physical association of *Drosophila* Tinagl1 with Wingless (Wg), a fly Wnt (Mulligan et al., 2012). In this interaction, Tinagl1 acted as a positive cofactor to stabilize and promote the diffusion of secreted Wg for its signaling beyond adjacent cells. In the zebrafish, Wnt/β-catenin signaling is required for ciliogenesis through the transcriptional activation of *foxj1a,* the master regulator of the motile cilia program (Caron et al., 2012). Experimentally, over-expression of the Wnt3a inhibitor Dkk or injection of morpholinos downregulating *wnt3a, β-catenin,* or *frizzled-10* disrupt ciliogenesis in the KV and pronephric ducts, resulting in heart looping defects and cystic distensions of the pronephric ducts similar to what we describe here for *tinagl1* knockdown. We note that Wnt3a signaling is also required for development of the pharyngeal arches, likely through multiple mechanisms (Sun et al., 2008). We previously found that *tinagl1* and *wnt3a* knockdowns result in similar defects in pharyngeal arch cartilages, and that a combination of sub-threshold doses of the TEx1 MO and Wnt3a MO, but not each MO separately with MM1 to give same final MO concentration, resulted in substantial losses of pharyngeal cartilages (Neiswender et al., 2017).

Here, we assayed pronephric duct cilia in an initial test of the hypothesis that Tinagl1 acts on cilia via Wnt/β-catenin signaling. Similar to the pharyngeal cartilage study (Neiswender et al., 2017), co-injection of sub-threshold doses of TEx1 MO and Wnt3a MO resulted in severe reduction of cilia length compared to normal length in uninjected embryos, while either mixed with MM1 to standardize the total MO dose did not show a reduction (**Supplemental Figure 5**). We also examined whether increasing *ctnnb2* expression could rescue pronephric cilia in the TEx1 MO embryos. The *ctnnb2* mRNA gave partial rescue of cilia length and number when coinjected with TEx1 MO (**Figure 6B, D, E**), similar in magnitude to what is seen for rescue by *tinagl1* mRNA. An *eGFP* mRNA co-injected at the same concentration had no rescuing effect (**Figure 6C**), supporting the likelihood that the *ctnnb2* and *tinagl1* mRNA rescue effects show specific roles for their gene products in formation of normal motile cilia. Together, the synergistic effect of Wnt3a and Tinagl1 MOs in severity of pronephric cilia defect, and the ability of the *ctnnb2* mRNA to partially rescue *tinagl1* morphant pronephric cilia, suggest that Tinagl1 may in fact exert effects on cilia through Wnt/β-catenin signaling during zebrafish embryo development.

### *tinagl1* knockdown results in spinal cord disorganization

In addition to the renal system, recall that we also observed high expression of *tinagl1* in the developing spinal cord (**Figure 1**). We hypothesized that this tissue would also be detrimentally affected by *tinagl1* knockdown. Therefore, we used immunofluorescence against acetylated tubulin to stain neurons and axon tracts in the spinal cord of 24-hpf embryos (**Figure 7**). We examined the presence of sensory neuron cell bodies dorsal to longitudinal axon tracts coursing in the anterior-posterior axis, and the outgrowth of CaP motor neuron axons along ventral somites (Lewis and Eisen, 2003). In a comparison of TEx1 MO- and MM1-injected embryos (**Figure 7A-D**), the MM1-injected embryos showed organization similar in quality to uninjected embryos. By contrast, the spinal cords in the *tinagl1* morphant embryos had reduced numbers of sensory neuron cell bodies and their longitudinal axon tracts appeared less organized.

**Figure 7.**
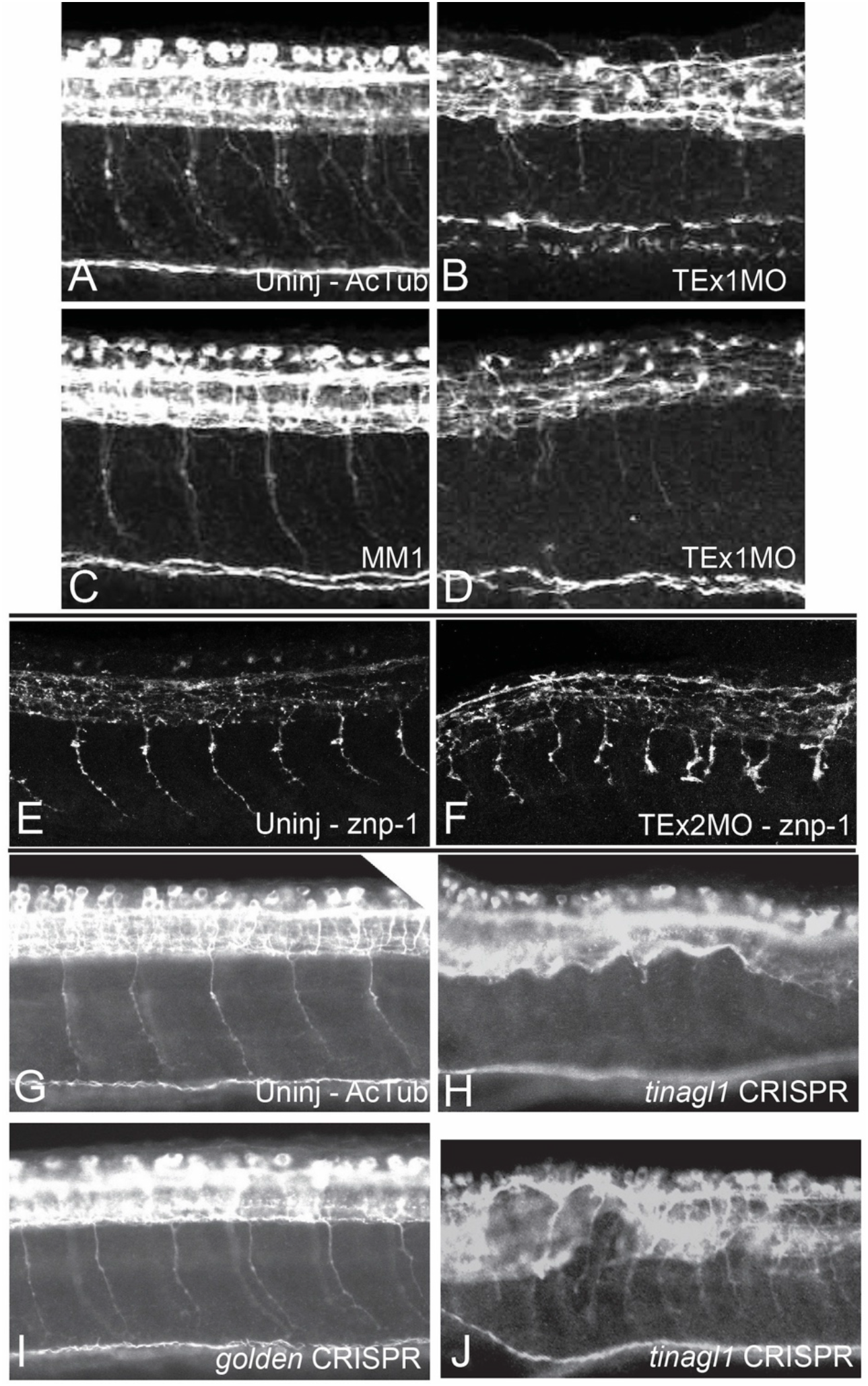
Spinal cord organization and motor axonal outgrowth are disrupted by acute *tinagl1* knockdown. (A-D) Anti-acetylated tubulin staining of TEx1 MO (B, D) *vs.* MM1 morphants (C) and uninjected siblings (A). (E-F) Znp-1 (synaptotagmin-2) monoclonal antibody staining highlights axonal branching and reduced outgrowth in TEx2 MO morphants (F) *vs.* uninjected siblings. (G-J) Anti-acetylated tubulin staining of *tinagl1* crispant (H, J) embryos *vs. golden* crispant (I) and uninjected siblings (G) similarly shows altered spinal cord tract architecture, reduced motor axon outgrowth, and reduced density of sensory neurons in *tinagl1* crispant but not *golden* crispant embryos.

We also detected reduced numbers of motor axons leaving the spinal cord in *tinagl1* morphants, as well as altered direction of their outgrowth with atypical branching. To confirm this observation, we used the znp-1 monoclonal antibody recognizing synaptotagmin-2 as an alternative marker for motor neurons (**Figure 7E, F**) (Trevarrow et al., 1990; Zeller et al., 2002). Staining confirmed reduced outgrowth and atypical branching of motor axons in *tinagl1* morphant embryos, as well as swellings in the tips of these axons that could indicate defects in axonal transport.

Somatic mosaic CRISPR-mediated knockdown of *tinagl1* also resulted in disruptions of spinal cord organization with reduction in motor axon outgrowth (**Figure 7G-J**). Use of *golden* sgRNA as a negative control did not show disruptions of spinal cord organization, indicating that these disruptions are not intrinsic to manipulation of embryos or to general CRISPR/Cas9 toxicity. While this nervous system analysis is qualitative, it suggests that there are requirements for *tinagl1* in spinal cord development, consistent with its transient expression along the spinal cord between 15-24 hpf.

## Discussion

Using two acute knockdown methods, we have demonstrated defects in embryonic tissues where Tinagl1 is expressed, supporting a conclusion that Tinagl1 is required for normal embryonic development in zebrafish. Somewhat surprisingly for an extracellular matrix protein characterized as being present predominantly in basement membranes, most of the observed defects could be best explained by a ciliopathy (Essner et al., 2005; Kramer-Zucker et al., 2005).

### Relationship to prior work on Tinagl1

Mainly in *in vitro* settings, mammalian Tinagl1 has shown significant and varied biological activities, including modulating integrin-based adhesion, stimulating angiogenesis, and inhibiting metastasis of an aggressive mouse mammary tumor cell line to lungs (Brown et al., 2010; Chen et al., 1996; Korpal et al., 2011; Li et al., 2007). This last finding is correlated to longer survival of breast cancer patients who have higher expression of *tinagl1* by their tumors. Furthermore, Tinagl1 has shown direct physical associations with Wnts, apparently increasing their activity and stability after secretion (Mulligan et al., 2012), and with the FN-binding integrin receptor and with EGF-R, blocking binding of FN and dimerization of EGF-R respectively (Shen et al., 2019). Despite these intriguing functional studies, a *tinagl1* knockout mouse line is viable and fertile and can be maintained over numerous generations (Takahashi et al., 2016), suggesting the protein is dispensable for mouse survival at least under un-challenged laboratory conditions. A *tinag* knockout mouse has not been described.

Some interesting factors may underlie the differences between the lack of obvious phenotypes in the knockout mouse and the activities demonstrated *in vitro* and in our zebrafish studies. These include genetic compensation by *tinag* or other ECM/signaling proteins when *tinagl1* expression is chronically abrogated in the knockout mouse (Rossi et al., 2015) and, for comparisons of whole animal work, examining post-natal versus embryonic roles. Notably, the knockout mouse line has fetal losses across pregnancy, though these were not characterized and were attributed to insufficiency of the placenta, a vascular tissue in which Tinagl1 is expressed (Tajiri et al., 2010; Takahashi et al., 2016). The single reported mouse developmental study, in which the kidney-enriched ortholog, Tinag, was acutely knocked down in mouse embryonic kidney explants, did show severe disruption of renal tubulogenesis and ureteric bud branching (Kanwar et al., 1999), suggesting either a critical role just for Tinag or that acute knockdown does not allow time for functional compensation by Tinagl1. Nonmammalian species have only one gene family member that in zebrafish shows greater aminoacid sequence similarity to mammalian Tinagl1, so would not be able to substitute its function by upregulating a more narrowly-expressed ortholog. In the future, we plan to generate and examine knockout *tinagl1* alleles in zebrafish, to further address and strengthen findings on the significance of this gene’s function in development and the signaling pathways with which it interacts *in vivo.*

### Potential roles of Tinagl1 in zebrafish embryonic development

The best-characterized functions for mammalian Tinagl1 and Tinag are in directly mediating integrin-based cell adhesion and in enhancing cell adhesion to major structural proteins of the basement membrane and interstitial matrix (Charonis et al., 1994; Chen et al., 1996; Kalfa et al., 1994; Li et al., 2007). These functions are consistent with Tinagl1 protein localization in mammals to the basement membrane of nearly all post-natal epithelia and vascular smooth muscle, and to interstitial matrices of the adrenal gland and renal glomerulus (Lennon et al., 2014; Li et al., 2007; Mukai et al., 2003; Wex et al., 2001). The nervous system defects we see in our zebrafish embryo models could readily be related to cell adhesion requirements in neuronal survival and for accurate axonal pathfinding and migration. Similarly, reduced neural crest cell populations within pharyngeal pouches could reflect decreased survival or migration related to cell adhesion defects.

Of other signaling pathways to which Tinagl1 has an attributed function, e.g., TGF-β, EGF-R, we have only looked at Wnt signaling, based on the biochemical evidence that fly Tinagl1 binds to a fly Wnt, Wg, with nanomolar affinity (Brown et al., 2010; Mulligan et al., 2012; Shen et al., 2019). The experimental approaches available in our acute knockdown models are not robust, but do suggest synergy of Wnt3a and Tinagl1 MOs in causing craniofacial cartilage and pronephric cilia defects, and that increasing β-catenin levels may partially ameliorate pronephric cilia defects in *tinagl1* morphants. These results could place Tinagl1 function as a positive cofactor within a model in which Wnt/β-catenin signaling activates transcription of *foxj1a* to establish the motile cilia program in zebrafish (Caron et al., 2012).

Regardless of specific mechanism, however, the observed defects in pronephric and KV cilia, along with their associated ciliopathy-like phenotypes (or phenocopies) of heart laterality defects, renal and pronephric duct cysts, and ventral body curvature, support a conclusion that Tinagl1 regulates ciliogenesis or cilia homeostasis (Essner et al., 2005; Kramer-Zucker et al., 2005). Tinagl1 would not be the first ECM protein to be associated with ciliopathy-like phenotypes (Zhang et al., 2020). Certain mutants of basement membrane laminin and a knockout of xylosyltransferase 2, an enzyme that initiates glycosaminoglycan synthesis, result in polycystic kidney disease through unknown mechanisms (Condac et al., 2007; Shannon et al., 2006; Vijaykumar et al., 2014). Mutations in the matrix metalloproteinase ADAMTS9 cause nephronophthisis-related ciliopathies in human patients, and endocytosed ADAMTS9 was surprisingly found to be required in periciliary vesicles for an early stage of ciliogenesis (Choi et al., 2019; Nandadasa et al., 2019). While Tinagl1 itself has not been linked to human or mouse ciliopathies, a single patient with pediatric nephronophthisis was found to have mutations in both alleles of the *tinag* gene (Sugimoto et al., 2011; Takemura et al., 2010).

Overall, we propose that zebrafish will provide a fruitful system in which to study the functional requirements for Tinagl1 in development, and to identify signaling pathways with which it interacts in a developmental context. Previously explored mainly in postnatal settings and in mice where an ortholog, Tinag, might be upregulated to compensate for loss of function, Tinagl1 appears to have important roles in development of the spinal cord, pronephros, and leftright patterning in zebrafish.

## Acknowledgments

We thank Jeffrey Mumm, Sammy Navarre, and David Kozlowski for getting us started in the zebrafish system; Scott Dougan and Wei-Chia Tseng for advice and protocols on doing CRISPR experiments at the start of 2015; Becky Burdine, Jonathan Eggenschwiler, other members of the UGA developmental biology community, Alberto Stolfi, Suneel Apte, and Thomas Stoeger for discussions; and Victoria L. Patterson for an insightful edit of the manuscript. Imaging was performed at the Transgenic Zebrafish and Cell Imaging Core Facilities at the Medical College of Georgia of Augusta University. Funding for this project came from the Pilot Study Research Program (Augusta University) and The Charles R. Silbereisen Fund of the Vanguard Charitable Gifts Program. EKL completed writing of the manuscript while supported by a Re-Entry to Biomedical Research Careers Supplement sponsored by NIAMS, and she thanks Becky Burdine and Sylvia B. Smith for making time and space for this work.

The authors declare no conflicts of interest.

