## Supplemental Data for "Acute knockdown of extracellular matrix protein Tinagl1 disrupts heart laterality and pronephric cilia in zebrafish embryonic development"

```
Zebrafish      MLRLWVLAAVSVLL----------------LLSEGGMTARTKRELAGPLHLRGIRDPFGS
Human          ---MWRCPLGLLLLLPLAG-----------HLALGAQQGRGRRELAPGLHLRGIRDAGGR
HTinag         ---MWTGYKILIFSYLTTEIWMEKQYLSQREVDLEAYFTRNHTVLQGTRFKRAI--FQGQ
                  :*      ::                  :   .   * :  *    . *.*    *

Zebrafish      YCQRRGGCCPGRNDQCTVPYL--DTICYCDLFCNRTVSDCCPDFWGHCLGTTP------P
Human          YCQEQDLCCRGRADDCALPYL--GAICYCDLFCNRTVSDCCPDFWDFCLGVPP------P
HTinag         YCR-NFGCCEDRDDGCVTEFYAANALCYCDKFCDRENSDCCPDYKSFCREEKEWPPHTQP
               **: .  ** .* * *.  :   .::**** **:*  ******: ..*           *

Zebrafish      YPPS-SCERNGHRFPSGSTYKENCNLCTCGQNGRWECEQHACLIEDDMIQEINRRDYGWR
Human          FPPIQGCMHGGRIYPVLGTYWDNCNRCTCQENRQWQCDQEPCLVDPDMIKAINQGNYGWQ
HTinag         WYP-EGCFKDGQHYEEGSVIKENCNSCTCSG-QQWKCSQHVCLVRSELIEQVNKGDYGWT
               : *  .* :.*: :   ..  :*** ***    :*:*.*. **:  ::*: :*: :***

Zebrafish      AANYSQFWGMTLDEGLRFRLGTKRPTRTIMNMNEMQMNMNGNDHLPSYFNAVDKWPGKIH
Human          AGNHSAFWGMTLDEGIRYRLGTIRPSSSVMNMHEIYTVLNPGEVLPTAFEASEKWPNLIH
HTinag         AQNYSQFWGMTLEDGFKFRLGTLPPSPMLLSMNEMTASLPATTDLPEFFVASYKWPGWTH
               * *:* ******::*:::****  *:  ::.*:*:   :     **  * *  ***.  *

Zebrafish      EPLDQGNCNASWAFSTAAVASDRISIQSMGHMTPQLSPQNLISCDTRHQDGCAGGRIDGA
Human          EPLDQGNCAGSWAFSTAAVASDRVSIHSLGHMTPVLSPQNLLSCDTHQQQGCRGGRLDGA
HTinag         GPLDQKNCAASWAFSTASVAADRIAIQSKGRYTANLSPQNLISCCAKNRHGCNSGSIDRA
                **** ** .*******:**:**::*:* *: *  ******:** ::::.** .* :* *

Zebrafish      WWFMRRRGVVTQDCYPFSPPEQSA-VEVARCMMQSRAVGRGKRQATAHCPNSHSYHNDIY
Human          WWFLRRRGVVSDHCYPFSGRERDEAGPAPPCMMHSRAMGRGKRQATAHCPNSYVNNNDIY
HTinag         WWYLRKRGLVSHACYPLFKDQN---ATNNGCAMASRSDGRGKRHATKPCPNNVEKSNRIY
               **::*:**:*:. ***:   :.        * * **: *****:**  ***.    * **

Zebrafish      QSTPPYRLSTNENEIMKEIMDNGPVQAIMEVHEDFFVYKSGIFRHTDVNYHKPSQYRKHA
Human          QVTPVYRLGSNDKEIMKELMENGPVQALMEVHEDFFLYKGGIYSHTPVSLGRPERYRRHG
HTinag         QCSPPYRVSSNETEIMKEIMQNGPVQAIMQVREDFFHYKTGIYRHVTSTNKESEKYRKLQ
               * :* **:.:*:.*****:*:******:*:*:**** ** **: *.  .  . .:**:

Zebrafish      THSVRITGWGEERDYSGRTRKYWIGANSWGKNWGEDGYFRIARGVNECDIETFVIGVWGR
Human          THSVKITGWGEETLPDGRTLKYWTAANSWGPAWGERGHFRIVRGVNECDIESFVLGVWGR
HTinag         THAVKLTGWGTLRGAQGQKEKFWIAANSWGKSWGENGYFRILRGVNESDIEKLIIAAWGQ
               **:*::****     .*:. *:* .*****  *** *:*** *****.***.:::..**:

Zebrafish      VTMEDMHNHHHHHGRRK
Human          VGMEDMGHH--------
HTinag         LTSSDEP----------
               :  .*
```

**Supplemental Figure 1. Protein sequence comparison of zebrafish Tinagl1 with human Tinagl1 and human Tinag.** Clustal 2.1 alignment (Sievers *et al.,* 2011) demonstrated the zebrafish sequence was 63.48% identical to the human Tinagl1 protein, and 50.33% identical to the human Tinag protein; the human Tinagl1 and Tinag shared 47.48% identity. A * below the protein sequences indicates 100% identity at that amino acid. Cysteines are highlighted in aqua, while red residues indicate amino acids of the pseudo-active cathepsin B-like site (S, H, N) where serine replaces cysteine. The navy blue bar overlies the predicted somatomedin B domain, while the black bar overlies the predicted lipocalin motif, which lies within the cathepsin B-like domain; domains and motif were predicted using the PROSITE database (Sigrist *et al.,* 2002; Sigrist *et al.,* 2013).

The roles of these domains in Tinagl1 function are unknown. For comparison from the literature, a well-characterized somatomedin B domain in vitronectin mediates cell adhesion via the transmembrane uPA receptor, with an overlapping PAI-1 binding site mediating cell detachment by competition (Blasi and Carmeliet, 2002; Seiffert and Loskutoff, 1991). A tandem pair of somatomedin B domains in autotaxin/NPP2 mediates binding to integrins (Hausmann *et al.*, 2011). The lipocalin motif found in Tinagl1 is conserved among kernal lipocalins as well as outlier proteins that like Tinagl1 lack 2 additional sequence motifs in an 8-stranded beta-barrel fold (Flower *et al.,* 1993). Mostly secreted proteins, the lipocalins bind hydrophobic molecules such as retinol and pheromones; the lipocalin motif in Tinagl1 has been suggested to bind the palmitate covalently attached to Wnt family members (Mulligan *et al.,* 2012). Tinag and Tinagl1 orthologs may be the only family of proteins in vertebrates that have an inactive cathepsin B-like domain, while secreted rat testin has an inactive cathepsin L-like domain and localizes to cell-cell junctions (Grima *et al.,* 1995). Trypanosomes and *Plasmodium* species have families of proteolytically-inactive cathepsin B-like secreted proteins with the same cysteine to serine alteration as seen in Tinagl1. These protozoan cathepsin B-like proteins have been postulated either to have a very narrow substrate range not detected with available reagents or to have alternative roles in diverting the immune system of their mammalian hosts; in either case, striking conservation in the binding pocket suggested that the protozoan proteins preserve ligand-binding properties and may have a peptidic partner (Mendoza-Palomares *et al*., 2008).

**A**

| | Curved body | Otolith defect (# or size) | Small eye | Jaw defect | Pericardial edema |
|---|---|---|---|---|---|
| **Uninject** 08/08/12 | 0% (0/25) | 0% (0/25) | 0% (0/25) | 0% (0/25) | 0% (0/25) |
| **MM1 5 ng** 08/08/12 | 0% (0/35) | 0% (0/35) | 0% (0/35) | 0% (0/35) | **13%** (2/16) |
| **TEx1MO 5 ng** 08/08/12 | **37%** (10/27) | **37%** (10/27) | **74%** (20/27) | **41%** (11/27) | **59%** (16/27) |
| **Uninject** 01/09/13 | 0% (0/25) | 0% (0/25) | 0% (0/25) | 0% (0/25) | 0% (0/25) |
| **Mismatch 10 ng** 01/09/13 | **13%** (2/16) | **6%** (1/16) | 0% (0/16) | 0% (0/16) | **13%** (2/16) |
| **TEx1 MO 10 ng** 01/09/13 | **87%** (27/31) | **68%** (21/31) | **81%** (25/31) | **100%** (31/31) | **97%** (30/31) |
| **P53/TEx1 15/10 ng** 01/09/13 | 100% (14/14) | 86% (12/14) | 100% (14/14) | 100% (14/14) | 100% (14/14) |

**B**

| Scored for: | Uninjected | MM 1 7.5 ng | TEx1MO 7.5 ng | Uninjected | MM2 20 ng | TEx2MO 20 ng |
|---|---|---|---|---|---|---|
| Curved body | 0% ( 0/33) | 0% (0/40) | **91%** (64/70) | 0% (0/58) | 0% (0/43) | **88%** (36/41) |
| Pericardial Edema | 0% ( 0/33) | 0% ( 0/40) | **94%** (66/70) | 0% (0/58) | 0% (0/43) | **85%** (35/41) |

**Supplemental Figure 2. Initial phenotypic assessment and titration of morpholinos.** We initially characterized defects that were visibly observable in live embryos, and scored several. These included: curved body – ventral body axis curvature; otolith defects (usually 1 but rarely 3 instead of 2 in one or both otic vesicles, though sometimes widely disparate sizes of 2 otoliths in same otic vesicle); small eyes, usually including optic fissure closure defects; jaw defects (described more fully in Neiswender *et al.,* 2017); and pericardial edema. Hydrocephalus and renal cysts were also often seen in the *tinagl1* morphants, but were not usually scored. (A) TEx1 MO gave dose-dependent defects, higher at 10 ng per embryo than at 5 ng per embryo, that were not or rarely seen with its MM1 mismatch control at same doses. Co-injection of a standard p53 MO did not rescue *tinagl1* morphants as might be expected if the defects were due principally to cell death. In fact, the defects were worsened, possibly a result of high total MO concentration when used at ratios seen in the literature; this led us not to use p53 MO as a routine control in subsequent experiments. (B) Final concentrations used for the majority of cilia-related experiments were 7.5 ng per embryo for TEx1 MO and MM1, and 20 ng/embryo for TEx2 MO and MM2. At these concentrations, TEx1 MO and TEx2 MO gave strong defects (all phenotypes), while the mismatch controls MM1 and MM2 gave rare or no defects.

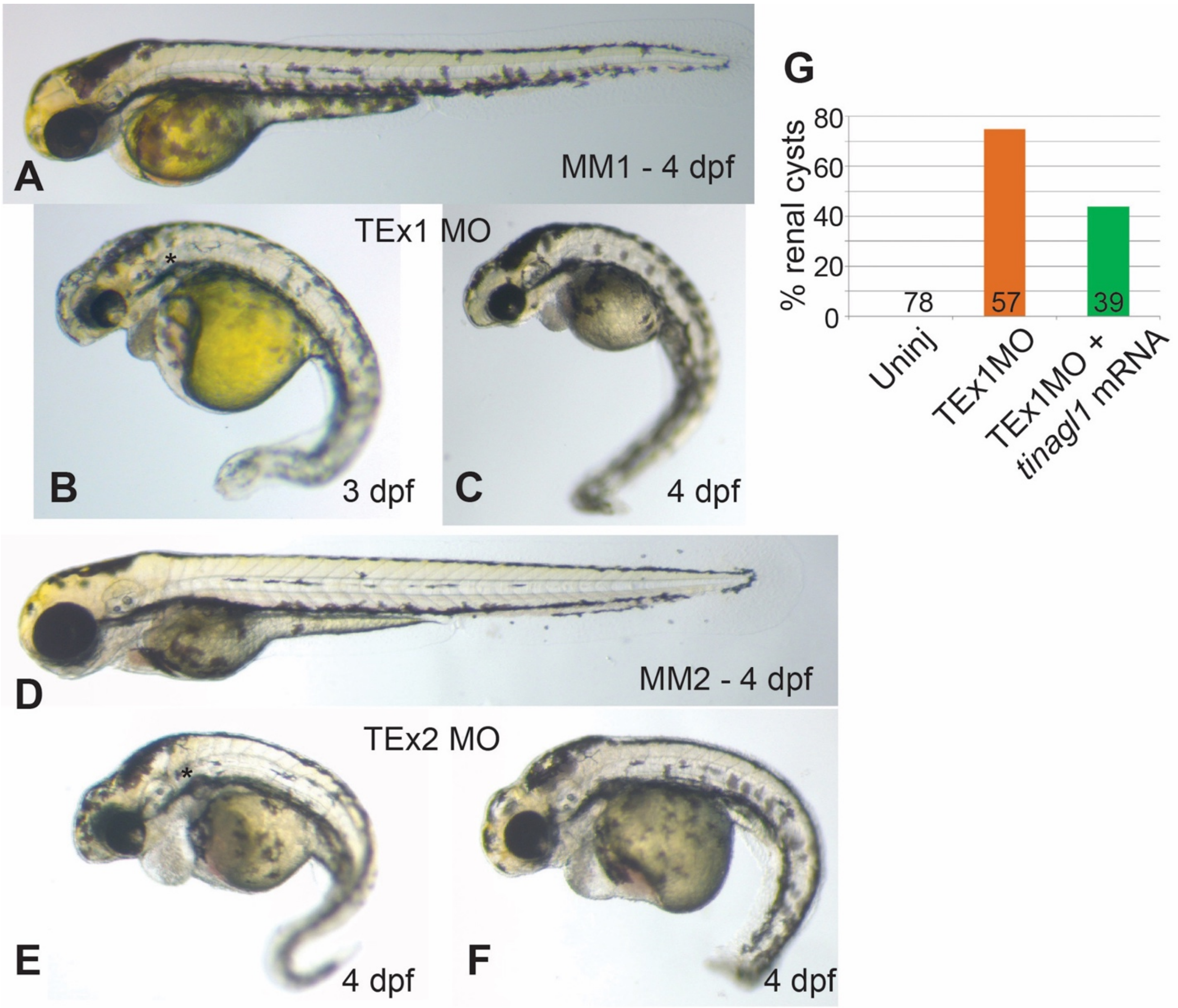


**Supplemental Figure 3. Comparison of MM and *tinagl1* MO phenotypes, and partial rescue of renal cysts by *tinagl1* mRNA.** (A-F) Typical appearance of 3-4 dpf embryos injected with 5 ng MM1 (A), 5 ng TEx1 MO (B, C), 20 ng MM2 (D), or 20 ng TEx2 MO (E, F). Renal cysts are marked by small asterisks in panels B and E. (G) Partial rescue of % embryos having renal cysts by co-injection of *tinagl1* mRNA with TEx1 MO.

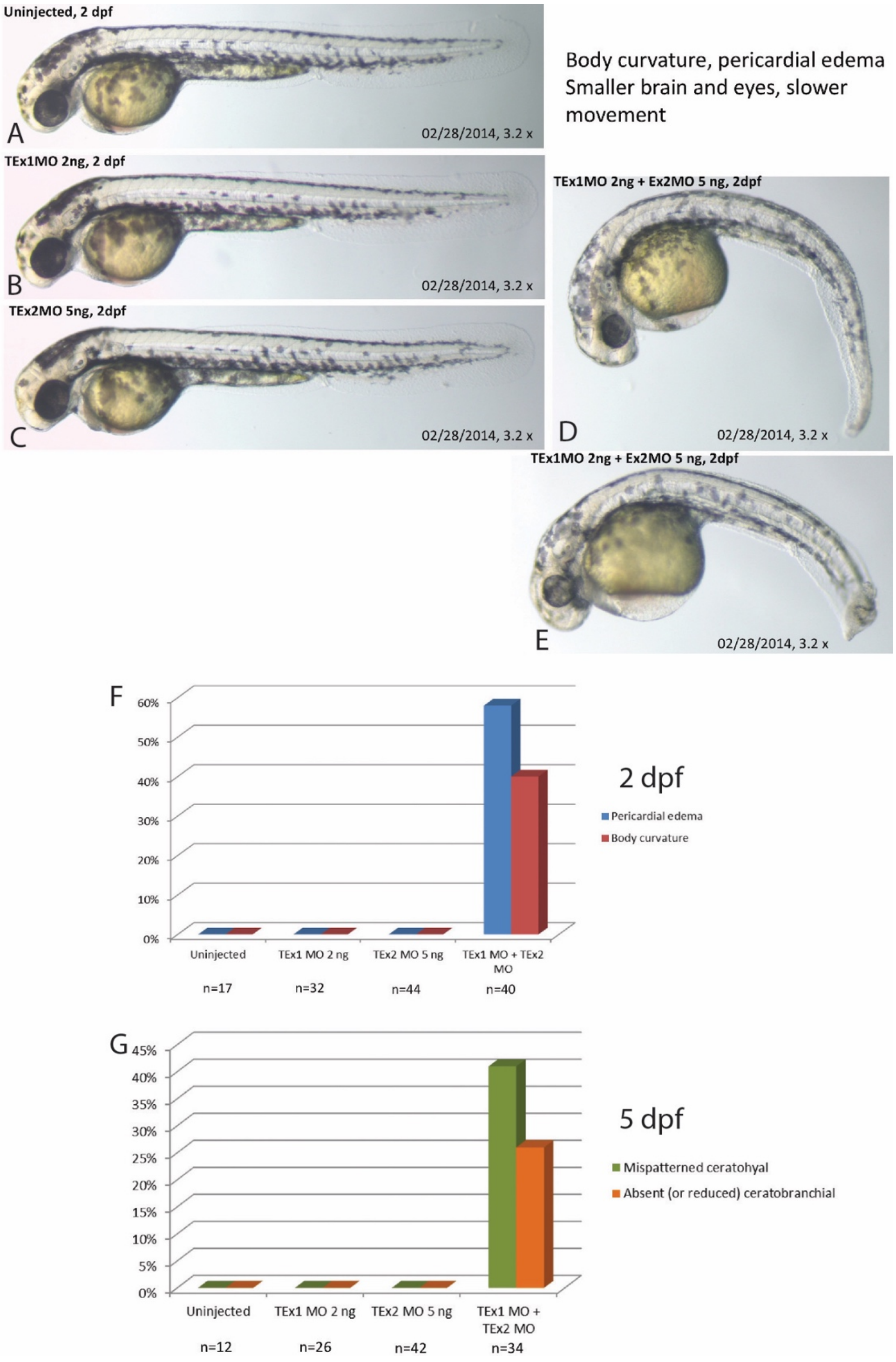


**Supplemental Figure 4. Combining sub-threshold doses of the TEx1 and TEx2 MOs causes the same constellation of defects as either MO alone at higher doses.** Embryos injected with 2 ng TEx1MO (B) or 5 ng TEx2MO (C) show normal morphology compared to uninjected embryos, while co-injection of 2 ng TEx1MO and 5 ng TEx2MO (D, E) resulted in defects similar in severity as described in Supplemental Figures 2B and 3. (F) Pericardial edema and ventral body curvature were scored for the indicated n of 2 dpf embryos in each treatment group. (G) Craniofacial defects (Neiswender *et al.,* 2017) were scored at 5 dpf.

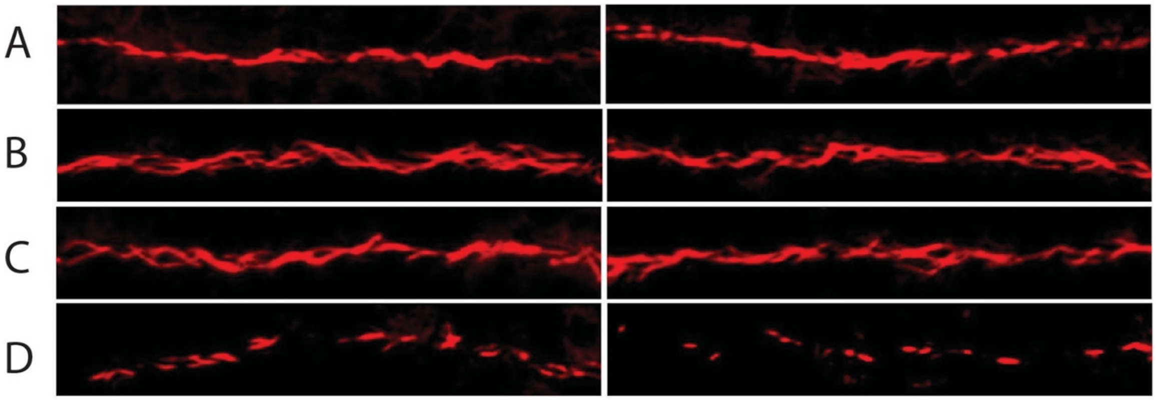

E

| *Pronephric cilia* | Uninjected (27 hpf) | TEx1 MO 3.5 ng MM 3.5 ng | Wnt3a MO 3.5 ng MM 3.5 ng | TEx1 MO 3.5 ng Wnt3a MO 3.5 ng |
|---|---|---|---|---|
| **Length (µm)** | 7.28 | 6.69 | 6.51 | 2.94 |
| STDEV | 0.70 | 0.97 | 0.98 | 0.68 |
| **Number per 100 µm** | 36 | 38 | 37 | 31 |
| STDEV | 1.7 | 4.5 | 4.5 | 1.5 |

**Supplemental Figure 5. Combined greater impact of low doses of Wnt3a MO and TEx1 MO on pronephric cilia.** (A-D) Two representative images of 24 hpf pronephric cilia for each condition including (A) uninjected, (B) TEx1 MO + MM1, each at 3.5 ng, (C) Wnt3a MO + MM1, each at 3.5 ng, and (D) TEx1 MO + Wnt3a MO, each at 3.5 ng. (E) Quantitation of cilia length and number per 100 µm for the four different conditions.

**References Unique to Supplemental Figure Legends**


Blasi, F. and Carmeliet, P. (2002) uPAR: a versatile signalling orchestrator. Nature Rev. Mol. Cell. Biol. 3: 932-943.

Flower, D. R., North, A. C. T., and Attwood, T. K. (1993) Structure and sequence relationships in the lipocalins and related proteins.

Grima, J., Zhu, L. J., Zong, S. D., Catterall, J. F., Bardin, C. W., Cheng, C. Y. (1995) Rat testin is a newly identified component of the junctional complexes in various tissues whose mRNA is predominantly expressed in the testis and ovary. Biol. Reprod. 52: 340-355.

Hausmann, J., Kamtekar, S., Christodoulou, E., et al. (2011) Structural basis of substrate discrimination and integrin binding by autotaxin. Nature Struct. Mol. Biol. 18: 198-205.

Mendoza-Palomares, C., Biteau, N., Giroud, C., Coustou, V., Coetzer, T., Authié, E., Boulangé, A., and Baltz, T. (2008) Molecular and biochemical characterization of a cathepsin B-like protease family unique to Trypanosoma congolense. Eukaryotic Cell 7: 684-697.

Seiffert, D. and Loskutoff, D. J. (1991) Evidence that type 1 plasminogen activator inhibitor binds to the somatomedin B domain of vitronectin. J. Biol. Chem. 266: 2824-2830.

Sievers, F., Wilm, A., Dineen, D., *et al.* (2011) Fast, scalable generation of high-quality protein multiple sequence alignments using Clustal Omega. Molec. Systems Biol. 7: 539.

Sigrist, C. J. A., Cerutti, L., Hulo, N., Gattiker, A., Falquet, L., Pagni, M., Bairoch, A., and Bucher, P. (2002) PROSITE: A documented database using patterns and profiles as motif descriptors. Briefings Bioinformatics 3: 265-274.

Sigrist, C. J. A., de Castro, E., Cerutti, L., Cuche, B. A., Hulo, N., Bridge, A., Bougueleret, L., and Xenarios, I. (2013) New and continuing developments at PROSITE. Nucl. Acids Res. 41:D344-347.
